# Requirement of the antimicrobial peptide CRAMP for macrophages to eliminate phagocytosed *E. coli* through an autophagy pathway

**DOI:** 10.1101/2020.07.23.218669

**Authors:** Keqiang Chen, Teizo Yoshimura, Wanghua Gong, Cuimeng Tian, Jiaqiang Huang, Giorgio Trinchier, Ji Ming Wang

## Abstract

Host-derived antimicrobial peptides play an important role in the defense against extracellular bacterial infections. However, the capacity of antimicrobial peptides derived from macrophages as potential antibacterial effectors against intracellular pathogens remains unknown. In this study, we report that normal (wild type, WT) mouse macrophages increased their expression of the cathelicidin-related antimicrobial peptide (CRAMP) after infection by viable *E. coli* or stimulation with inactivated *E. coli* and its product LPS, a process involving activation of NF-κB followed by protease-dependent conversion of CRAMP from an inactive precursor to an active form. The active CRAMP was required by WT macrophages to eliminate phagocytosed *E. coli*, with participation of autophagy-related proteins ATG5, LC3-II, and LAMP-1 as well as conjugation of the bacteria with p62. The autophagy-mediated elimination of *E. coli* was impaired in *CRAMP*^*−/−*^ macrophages resulting in retention of intracellular bacteria and fragmentation of macrophages. These results indicate CRAMP as a critical component in autophagy-mediated clearance of intracellular *E. coli* by macrophages.

## Introduction

Macrophages comprise an essential part of the innate immune system in response to bacterial infections **(Rosenberger and Finlay, 2003)**. Because macrophages are highly phagocytic and are readily confronted by pathogenic bacteria, they must be equipped with effective mechanisms either for killing bacteria or controlling their replication to avoid becoming a reservoir of infection. For example, colon macrophages residing in the subepithelial lamina propria (LP) represent the first line defense against invading pathogens hence act as crucial sentinels for the maintenance of colon homeostasis **(Mowat and Agace, 2014)**. *E. coli* belongs to the family of *Enterobacteriaceae, phylum* of *Proteobacteria,* which although constitutes a minor fraction of the microbiome found in human gastrointestinal tract **(Bailey et al., 2010)**, is the most common cause of intestinal and extra-intestinal diseases **(Conway and Cohen, 2015; Foster, 2004; Katouli, 2010)**. Many host factors including inflammation and genetic predisposition markedly alter the colonic microbial composition and support the growth of either resident or introduced aerobic bacteria, particularly those of the *Enterobacteriaceae* family **(Lupp et al., 2007)** such as *E. coli,* which are elevated in inflammatory bowel diseases (IBD) **(Bambou et al., 2004; Martin et al., 2004; Rhodes, 2007; Zhang et al., 2017)**. *E. coli* species are also increased in colorectal cancer tissues in association with DNA damage in epithelial cells **(Arthur et al., 2012; Dejea et al., 2018)**. Previous studies showed that increased mucosal adherent *E. coli* plays a central role in the pathogenesis of human IBD and colon cancer **(Martin et al., 2004)**. Despite the observations that macrophages are capable of controlling *E. coli* expansion by direct killing or phagocytosis, there is scarce data concerning the mechanisms by which macrophages eliminate phagocytosed *E. coli.*

Autophagy is utilized by macrophages to eliminate intracellular or phagocytosed bacteria **(Deretic, 2011; Levine et al., 2011) t** o exert a housekeeping function therefore plays a protective role in maintaining cellular homeostasis **(Moreau et al., 2010).** Autophagy process in macrophages is activated in response to many stress conditions including starvation, endoplasmic reticulum disfunction, oxidative damage, and exposure to chemicals, radiation and hypoxia **(Mizushima and Komatsu, 2011)**. Bacterial infection and inflammation are also able to trigger autophagy in macrophages and other immune cells **(Saitoh and Akira, 2010)**. When activated in infected macrophages, autophagy promotes the clearance of pathogenic bacteria including *Salmonella typhimurium, Shigella flexneri* **(Deretic and Levine, 2009)** and *Mycobacterium tuberculosis* **(Rekha et al., 2015)**. Bacteria initiate autophagy in macrophages mainly via their pathogen-associated molecular patterns (PAMPs) and damage-associated molecular patterns (DAMPs). Cell surface recognition and cytosolic sensing of invading pathogens by these molecules result in signaling cascades that promote rapid and localized autophagy machinery assembly. For instance, as a cytosolic sensor in macrophages, c-GAS recognizes bacterial DNA to trigger autophagy activation, resulting in ubiquitination of the bacterium or its phagosome by ubiquitin ligases Parkin and Smurf1. Ubiquitin chains subsequently bind to autophagy adaptors, such as p62 and NDP52 that recruit LC3 to deliver bacteria into an autophagosome. In addition, damaged phagosome is also targeted by autophagy via the recognition of host glycan present in the phagosomal lumen through cytosolic lectins of the galectin family. The process is tightly regulated by more than 30 autophagy-related gene products (ATGs). Upon autophagy activation, ATGs, serine/threonine kinase ULK1, and Beclin-1, in association with Atg14 and type III phosphatidylinositol 3-kinase (PI3K) Vps34, promote the formation of a cup-shaped isolation membrane to engulf the cargo to form a double-membrane autophagosome, which then fuses with lysosomes to form an autolysosome in which the engulfed cargo is degraded **(Klionsky, 2010)**. However, the role of autophagy process in macrophage elimination of phagocytosed *E. coli* is unclear.

LL-37 in human and its mouse orthologue CRAMP are cathelicidin-related antimicrobial peptides, which belong to a family of host-derived antibacterial polypeptides **(Zhang et al., 2019)**. LL-37 and CRAMP are amphipathic *a*-helical peptides that bind to negatively charged groups of the bacterial outer membrane causing disruption of the cell wall **(Scott and Hancock, 2000)**. In mouse macrophages, CRAMP is upregulated by infection of intracellular pathogens such as *Salmonella typhimurium* **(Rosenberger et al., 2004)** or *M. smegmatis* **(Sonawane et al., 2011).** CRAMP is an essential component in host anti-microbial defense, by directly impairing the replication therefore killing intracellular pathogens **(Rosenberger et al., 2004; Sonawane et al., 2011)** by macrophages as well as participating in autophagy process to eliminate bacteria. In human macrophages, LL-37 is not only directly bactericidal but also serves as a mediator of vitamin D3-induced autophagy to activate the transcription of autophagy-related genes Beclin-1 and Atg5, therefore indirectly participating in the elimination of intracellular bacteria **(Yuk et al., 2009)**. However, it is not known whether CRAMP in mouse macrophages acts as a part of an antibacterial effector mechanism against phagocytosed *E. coli*.

In this study, we demonstrate that CRAMP is involved in the elimination of phagocytosed *E. coli* by mouse macrophages as shown by the retention of phagocytosed *E. coli* of macrophages deficient in CRAMP. We further provide evidence that CRAMP deficiency results in reduced expression of autophagy-related molecules ATG5, LC3-II, LAMP-1 and p62 and impaired degradation of *E. coli* conjugated with p62 in macrophages.

## Results

### Stimulated of CRAMP production in macrophages by *E. coli* products

To obtain evidence for the importance of CRAMP for macrophages to eliminate phagocytosed *E. coli*, we generated macrophages from bone marrow (BM) cells of control *CRAMP*^*+/+*^ mice (*LysMCre^−^CRAMP^F/F^*). After infection with *E. coli* isolated from the feces of naïve mice, the production of CRAMP by macrophages progressively increased and reached the maximal level by 20 h (**Fig. 1A, B**). Inactivated *E. coli* also stimulated control macrophages to produce CRAMP as confirmed by Western blotting (**Fig. 1C**).

**Figure 1.**
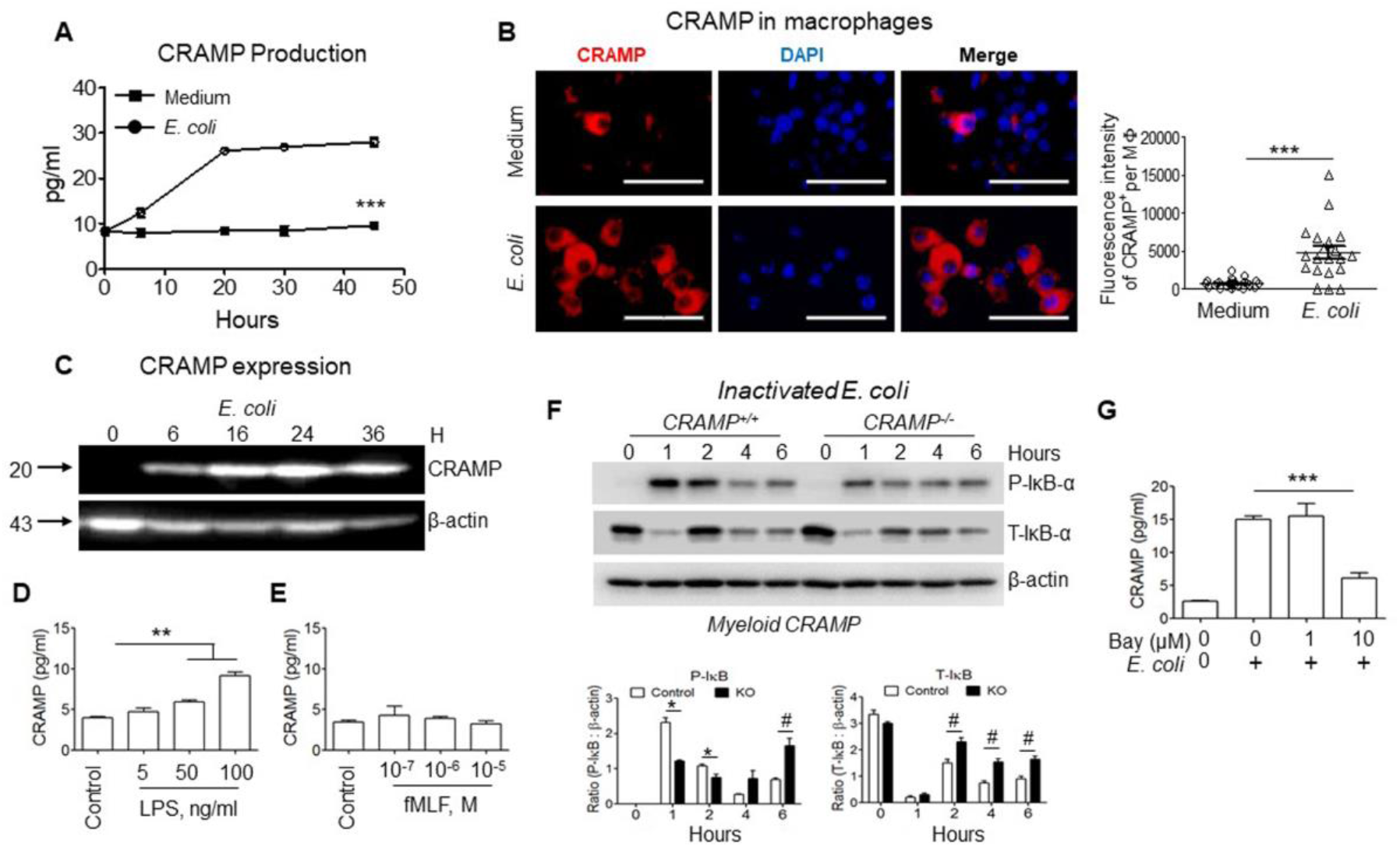
CRAMP production induced by *E. coli* products in macrophages. **A.** Production of CRAMP by macrophage. macrophages from BM of control (*LysMCre*^*−*^*CRAMP*) mice were seeded in 96 well plates at 1.5 × 10^5^/well and infected with *E. coli* O22H8 (20 μl/well, at 6.0× 10^8^ CFU/ml). Culture supernatants were harvested at 0, 6, 20, 30, and 45 h for measurement of CRAMP by ELISA, n = 3/group. \*\*\**P* < 0.001, significantly increased CRAMP in supernatants of *E. coli* infected cells compared to cells treated with medium alone at 20, 30, and 45 h (two-tailed Student’s t test) **B.** Detection of increased CRAMP in macrophages infected by *E. coli* O22H8. Red: CRAMP; Blue: DAPI. Scale bar = 30 μm (**Left panels**). **Right panel**: Quantitation of CRAMP positive staining spots per macrophage. Shown is the immunofluorescence intensity per macrophage, n = 10-16 fields/group. \*\*\**P* < 0.001 by two-tailed Student’s t test. **C.** Upregulation of CRAMP in macrophages stimulated with inactivated *E. coli* O22H8. macrophages from BM of control mice were stimulated with inactivated *E. coli* O22H8 (100 μl at 6 × 10^8^ CFU/ml) at 37°C then were lysed at the indicated time points. The cell lysates were measured for CRAMP by western blotting. **D-E.** LPS or fMLF stimulated CRAMP production by macrophages. macrophages from BM of control (*LysMCre*^*−*^*CRAMP*) mice were seeded in 96 well plates at 1.5 × 10^5^/well and stimulated with LPS **(D)** or fMLF **(E)** at indicated concentration for 24 h. The culture supernatants were measured for CRAMP by ELISA, n = 3/group. \*\**P* < 0.01 by 1-way ANOVA with Kruskal-Wallis Test. **F.** Reduced IκB activation in CRAMP^−/−^ macrophages from BM of *Myeloid CRAMP*^*−/−*^ (KO) mice by inactivated *E. coli* O22H8. **Lower panels**: Densitometry measurement of the ratio of phosphor-IκB against β-actin (**left**) and ratio of total-IκB: β-actin (**right**). \**P* < 0.05, #*P* < 0.05, by two-tailed Student’s t test. **G.** IκB inhibitor BAY117082 attenuated CRAMP production of macrophage. Macrophages were seeded in 96 well plate at 1.5 × 10^5^/well and cultured in the presence of different concentrations of BAY117082 for 1 h at 37°C before stimulation with inactivated *E. coli* O22H8 (20 μl at 6 × 10^8^ CFU/ml) for an additional 20 h. The supernatant was harvested for measurement of CRAMP by ELISA. \*\*\**P* < 0.001, 1-way ANOVA with Kruskal-Wallis Test.

In addition, Lipopolysaccharide (LPS) as the principal component of Gram-negative bacteria such as *E. coli* **(Raetz and Whitfield, 2002)** stimulated control macrophages to produce CRAMP in a dose-dependent manner **(Fig. 1D)**. In contrast, another product of *E. coli*, the chemotactic peptide N-formyl-methionyl-leucyl-phenylalanine (fMLF) **(Schiffmann et al., 1975)**, failed to stimulate macrophages to produce CRAMP (**Fig. 1E**).

We further observed revealed that stimulation of control macrophages by inactivated *E. coli* induced rapid phosphorylation of IκB, shown by an increase in total IκB due to *de novo* synthesis (**Fig. 1F**) **(Karin, 1999)**. In contrast to control macrophages, there was a significantly diminished phosphorylation of IκB-α in *CRAMP*^*−/−*^ macrophages from *Myeloid CRAMP*^*−/−*^ mice (**Fig. 1F**). The CRAMP production by control macrophages in response to inactivated *E. coli* was attenuated by a selective IκB-α inhibitor BAY117082 (**Fig. 1G**). Thus, activation of NF-κB is critical for macrophages to produce CRAMP in response to stimulation by *E. coli* and its product LPS.

### Requirement of CRAMP for macrophages to eliminate phagocytosed *E. coli*

To examine the role of CRAMP in macrophage elimination of phagocytosed *E. coli*, a mouse RAW 264.7 cell line, used as an *in vitro* model, was co-cultured with inactivated *E. coli* for 20 h. RAW 264.7 cells expressed high level of CRAMP with few endocytosed inactivated *E. coli* (**Fig. 2A, Upper panel**). Preincubation of RAW 264.7 cells with BAY117082 reduced the production of CRAMP with increased phagocytosed *E. coli* within the cells (**Fig. 2A, Lower panel**). The bactericidal activity of CRAMP was also shown by a synthetic peptide that directly killed *E. coli in vitro* (**Fig. 2B**).

**Figure 2.**
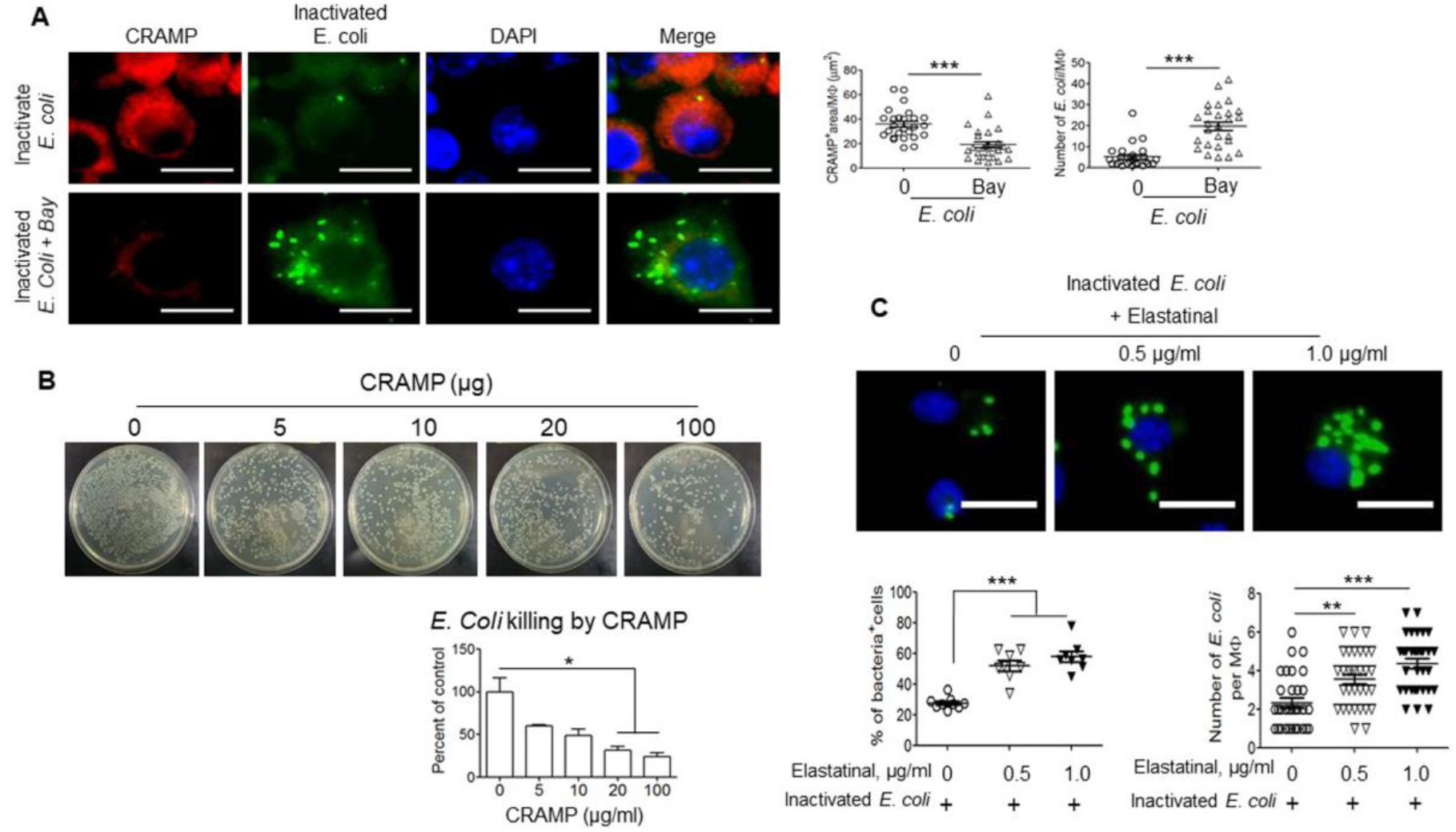
Requirement of CRAMP for macrophages to eliminate phagocytosed *E. coli*. **A.** Reduction in CRAMP production and degradation of inactivated *E. coli* O22H8 in macrophages treated with BAY117082. **Left panels:** CRAMP^+/+^ macrophages from BM of control mice were seeded in 35 mm dishes with 14 mm coverslips at 1× 10^6^/dish. The cells were then cultured in the presence or absence of 10 μm Bay117082 for 1 h at 37°C before stimulation with inactivated *FITC* labeled *E. coli* O22H8 (100 μl of 6 × 10^8^ CFU/ml) for an additional 20 h. The cells were then stained with an anti-mouse CRAMP Ab. Red: CRAMP; Green: inactivated *E. coli*-FITC; Blue: DAPI. Scale bar= 10 μm. **Right panels**: **Left side**: Reduced CRAMP production in CRAMP^+/+^ macrophages treated by Bay117082. **Right side**: Delayed elimination of inactivated *E. coli* by macrophages treated with Bay117082, \*\*\**P* < 0.001, two-tailed Student’s t test. **B.** Killing of *E. coli* O22H8 by synthetic CRAMP. *E. coli* O22H8 was diluted to a concentration of 5 × 10^4^ in 100 μl/well in 96-well plates. Various concentrations of synthetic CRAMP were added to the culture for incubation at 37°C for 2 h. The bacteria cultured with or without CRAMP were serially diluted at 1:5 with sterile PBS and plated on LB agar in triplicates to examine colony formation. **Lower panel:** Quantitation of effect of CRAMP at different concentration to kill *E. coli*. \**P* < 0.05, by 1-way ANOVA with Kruskal-Wallis Test. **C.** Reduction of degradation of intracellular inactivated *E. coli* O22H8 by macrophages in the presence of an elastase inhibitor Elastatinal. Upper panels: Murine Raw 264.7 cells seeded in 35 mm dishes with 14 mm coverslips at 1× 10^6^/dish were cultured in the absence or presence of Elastatinal (0.5 or 1 μg/ml) for 1 h at 37°C before stimulation with inactivated *E. coli-FITC* (100 μl at 1 × 10^8^ CFU/ml) for an additional 20 h. Green: inactivated *E. coli*-FITC; Blue: DAPI. Scale bar = 10 μm. Lower left panel: Quantitation of the cells positive with bacteria (%). Lower right panel: Quantitation of bacteria number per Raw 264.7 cell. The experiment was repeated three times, n = 8 fields/group. \*\**P* < 0.01, \*\*\**P* < 0.001, 1-way ANOVA with Kruskal-Wallis Test.

CRAMP is normally stored in lysosomes of macrophages as an inactive precursor, which is converted to an active form through cleavage by proteases **(Shinnar et al., 2003; Zanetti, 2004)** such as intracellular elastase-like serine protease **(Rosenberger et al., 2004)**. We found that Elastatinal, an elastase inhibitor, attenuated the capacity of macrophages to eliminate phagocytosed *E. coli* (**Fig. 2C**). Therefore, CRAMP production and conversion are critical for macrophages to eliminate both phagocytosed and extracellular *E. coli*.

### Reduced capacity of *CRAMP*^*−/−*^ macrophages to eliminate phagocytosed *E. coli* (K, I moved the source of CARMP+ macrophages to preceding paragraphs)

As shown in **Fig. 3A**, after infection with *E. coli*, the number of the bacteria was significantly increased in *CRAMP*^*−/−*^ macrophages (from *LysMCre*^*+*^*CRAMP*^*F/F*^ myeloid conditional *CRAMP*^*−/−*^ mice) as compared with *CRAMP*^*+/+*^ control macrophages 4 h after infection. By 20 h, many *CRAMP*^*−/−*^ macrophages disintegrated that allowed the formation of numerous extracellular bacterial colonies. By contrast, only a small number of bacteria were visible in macrophages from control macrophages.

**Figure 3.**
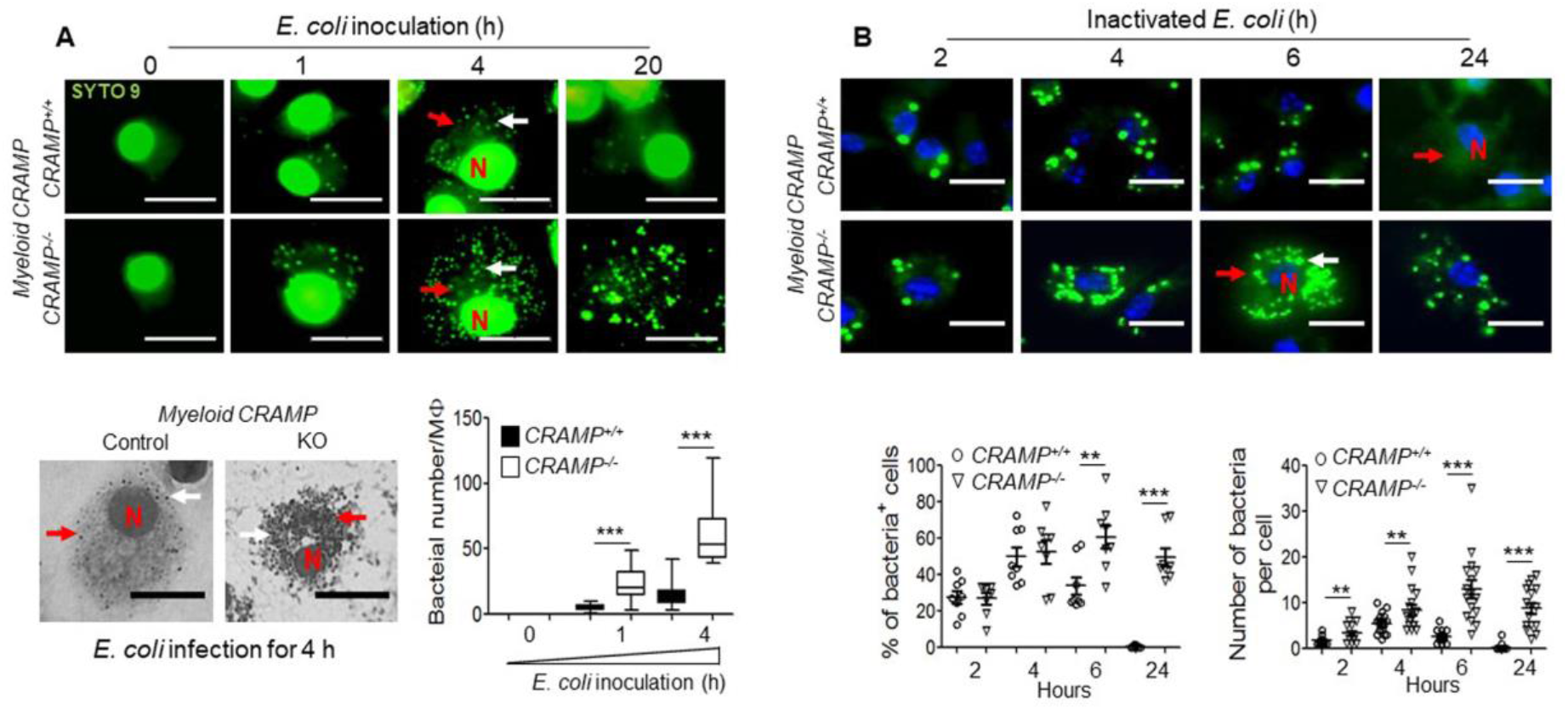
Attenuation of the capacity of CRAMP^−/−^ macrophages to eliminate phagocytosed *E. coli*. **A.** Killing of phagocytosed *E coli* O22H8 by macrophages. CRAMP^+/+^macrophages from BM of control and CRAMP^−/−^ macrophages from BM of *Myeloid CRAMP*^*−/−*^ mice were seeded in 35 mm dishes with 14 mm coverslips at 1× 10^6^/dish. The cells were then infected with *E. coli* O22H8 (100 μl at 6.0 × 10^8^ CFU/ml) and harvested at indicated time points for staining with SYTO 9 to reveal intracellular bacteria. N: nuclei, White arrows: *E. coli*, Red arrow: cell membrane. Scale bar = 10 μm. **Lower left panel:** Black pictures showed phagocytosed *E. coli* in macrophages for 4 h, Scale bar = 10 μm, N: nuclei, Red arrows: *E. coli*. **Lower right panel**: Quantitation of *E. coli* in each macrophage. \*\*\**P* < 0.001, significantly higher number of *E. coli* O22H8 in CRAMP^−/−^ macrophages (1-way ANOVA with Kruskal-Wallis Test). **B.** Failure of CRAMP deficient macrophages to eliminate phagocytosed inactivated *E. coli* O22H8. **Upper panels:** CRAMP^+/+^ and CRAMP^−/−^ macrophages were seeded in 35 mm dishes with 14 mm coverslips at 1× 10^6^/dish. The cells were stimulated with *FITC* labeled inactivated *E. coli* O22H8 (100 μl at 1 × 10^8^ CFU/ml) then harvested at indicated time points. Green: Inactivated *E. coli* O22H8, Blue: DAPI. N: nuclei, White arrows: *E. coli*, Red arrow: cell membrane. Scale bar = 10 μm. **Lower left panel:** Quantitation of macrophages (%) phagocytosd inactivated *E. coli* O22H8. **Lower right panel:** Quantitation of phagocytosed inactivated *E. coli* O22H8 in single macrophage. The experiment was repeated three times, n = 7-12 fields/group. \*\**P* < 0.01; \*\*\**P* < 0.001, 1-way ANOVA with Kruskal-Wallis Test.

In addition, in *CRAMP*^*+/+*^ control macrophages, the percent of the cells phagocytosing *E. coli* and the number of *E. coli* per cell reached a peak at 4 h, followed by a reduction at 6 h, with only very few bacteria visible at 24 h **(Fig. 3B, Upper panel)**. In contrast, in *CRAMP*^−/−^ macrophages, the percent of the cells phagocytosing *E. coli* and the number of *E. coli* per cell reached a peak at 6 h and a considerable number of bacteria remained in the cells at 24 h **(Fig. 3B, Lower panel)**. These results indicate that CRAMP was required for macrophages to timely eliminate phagocytosed *E. coli*.

### Involvement of autophagy pathway in CRAMP-mediated elimination of phagocytosed *E. coli* by macrophages

We then tested whether lysosomal hydrolases in macrophages are required for autophagic elimination of inactivated *E. coli*. Treatment of *CRAMP*^*+/+*^ control macrophages or RAW264.7 cells with E64d, an inhibitor of cathepsins B and L, or pepstatin A, an inhibitor of cathepsin D, that suppress autolysosomal digestion, protected *E. coli* from autophagic elimination by the cells **(Fig. 4A-C)**. Thus, lysosomal proteases are important for autophagic degradation of inactivated *E. coli* by macrophages, indicating the dependence on autophagy.

**Figure 4.**
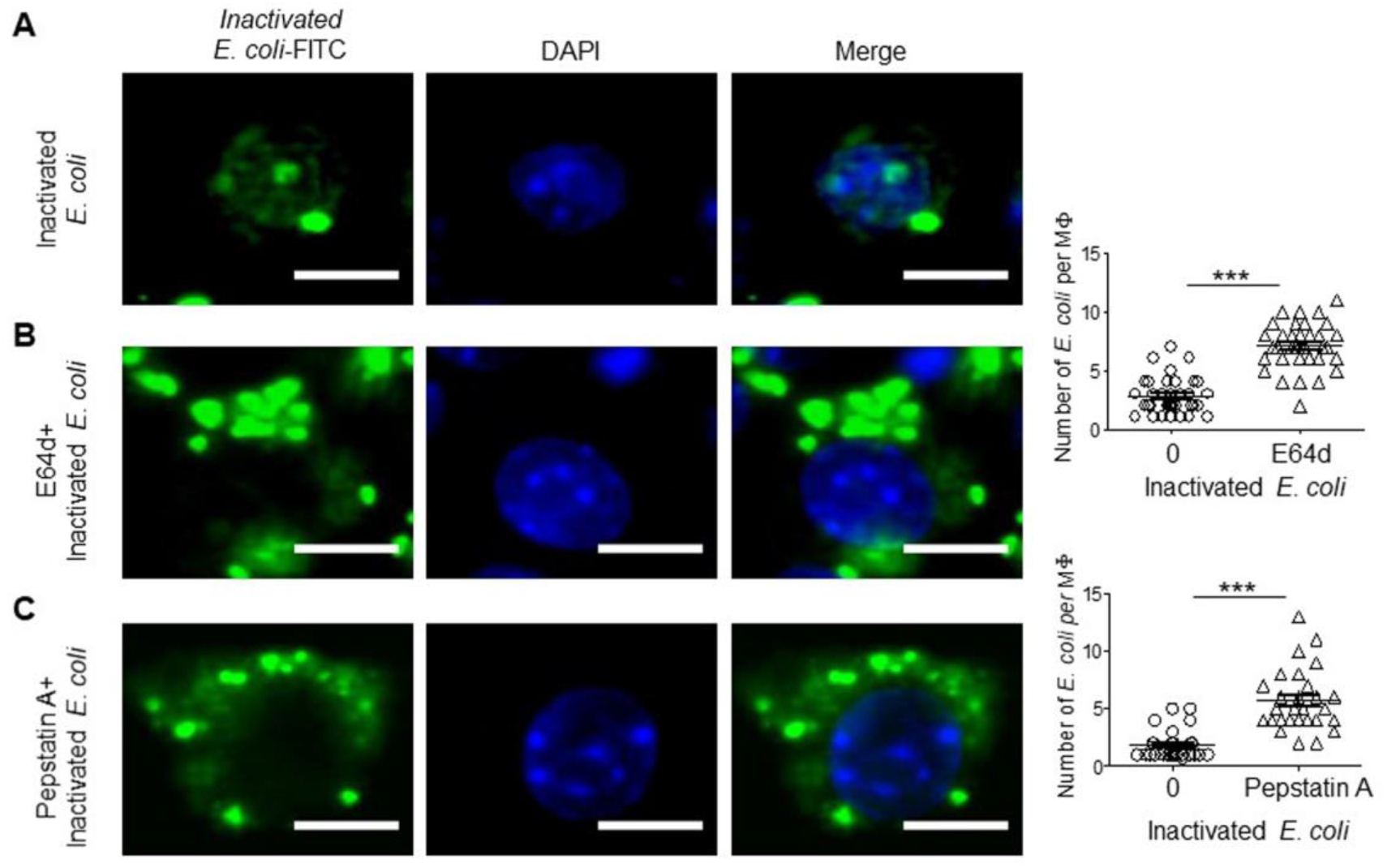
Delayed elimination of inactivated *E. coli* O22H8 in macrophages by inhibitors of auto-phagolysosomes. CRAMP^+/+^ macrophages were pretreated with E64d (1μg/ml) or Pepstatin A (10 μg/ml) for 1 h at 37°C before stimulation with FITC labeled inactivated *E. coli* O22H8 (100 μl at 6 × 10^8^ CFU/ml) for an additional 20 h. **A.** Control group, **B.** E64d treatment group, **C.** Pepstatin treatment group. Green: inactivated *E. coli*-FITC; Blue: DAPI. Scale bar = 5 μm. **Right upper panel**: Delayed elimination of phagocytosed inactivated *E. coli* from E64D treatment group. **Right lower panel**: Delayed elimination of inactivated *E. coli* from Pepstatin A treatment group. \*\*\**P* < 0.001, two-tailed Student’s t test.

We further found that there was a reduced expression of the autophagy-related protein ATG5, which is involved in the extension of the phagophoric membrane in autophagic vesicles **(Matsushita et al., 2007)** in *CRAMP*^−/−^ macrophages as compared to *CRAMP*^*+/+*^ control counterparts (**Fig. 5A)**. Under normal conditions, ATG5 forms complexes with ATG12 and ATG16L1, necessary for the conjugation of LC3-I (microtubule-associated proteins 1A/1B light chain 3B) to phosphatidylethanolamine (PE) to form LC3-II **(Otomo et al., 2013)**. However, LC3-II formation was also reduced in *CRAMP*^*−/−*^ macrophages after phagocytosis of inactivated *E. coli* (**Fig. 5A**). The adaptor protein p62 is an autophagy-targeting molecule recognizing ubiquitinated cytoplasmic components and delivering them for degradation **(Ponpuak et al., 2010)**. *CRAMP*^*−/−*^ macrophages showed reduced expression of p62 at 2 and 4 h, as compared to *CRAMP*^*+/+*^ control macrophages after stimulation with inactivated *E. coli* **(Fig. 5B)**. However, intracellular p62 accumulation was similar between *CRAMP*^*−/−*^ and *CRAMP*^*+/+*^ control macrophages at 6 h after *E. coli* stimulation **(Fig. 5B)**, indicating that the production and degradation of p62 induced by *E. coli* was impaired in *CRAMP*^*−/−*^ macrophages.

**Figure 5.**
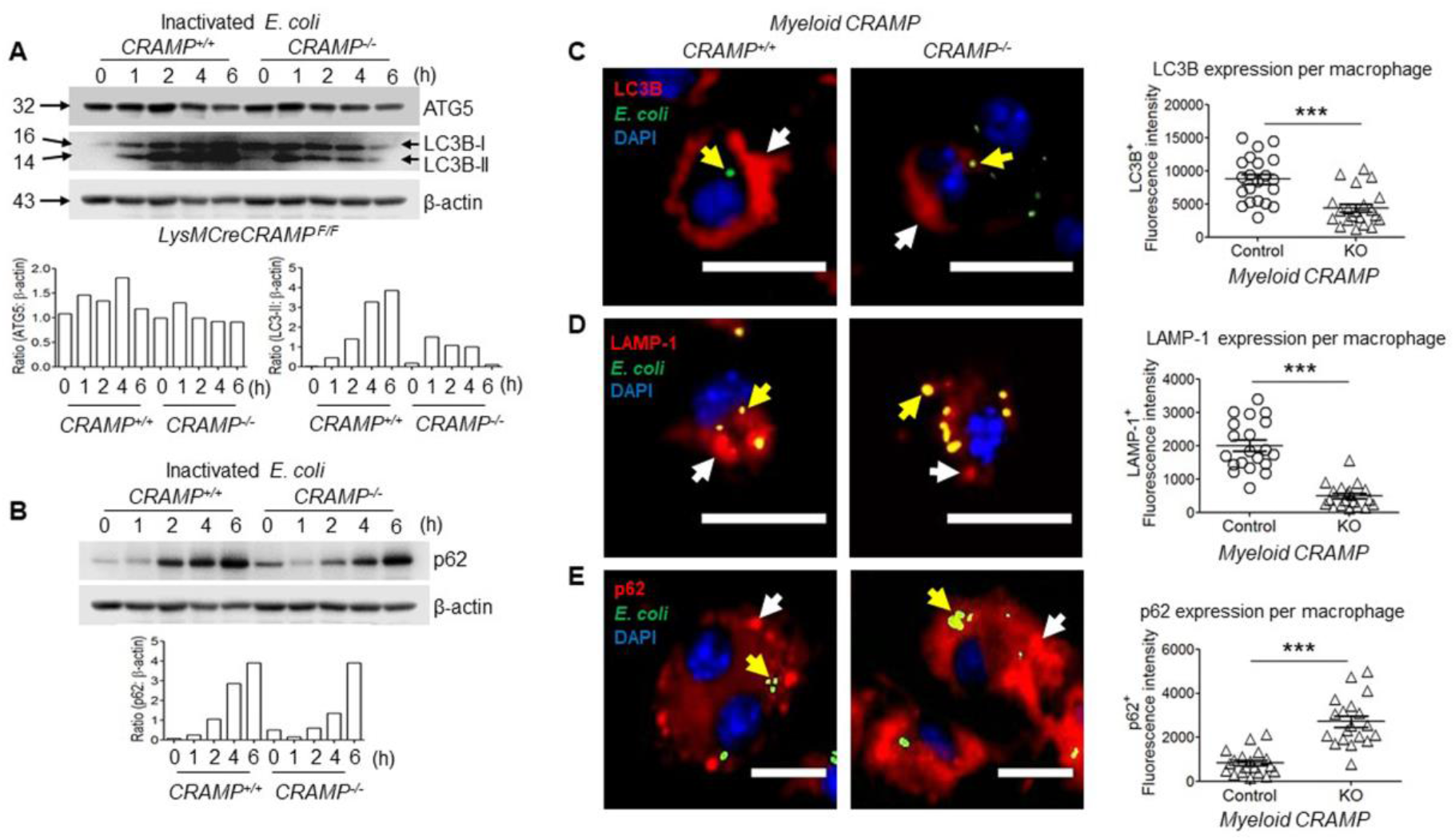
Involvement of autophagy pathway in CRAMP-mediated elimination of inactivated *E. coli* by macrophages. **A.** Activation of autophagy-related proteins ATG5 and LC3B-II in macrophages. CRAMP^+/+^ and CRAMP^−/−^ macrophages were cultured in the presence of inactivated *E. coli* O22H8 (100 μl at 6 × 10^8^ CFU/ml) at 37°C then lysed at the indicated time points. The cell lysates were measured for ATG5, LC3-I and LC3-II proteins by Western blotting. **Lower panels**: Ratio of ATG5: β-actin (**left**), Ratio of LC3-II: β-actin (**right**). **B.** Activation of autophagy-related proteins p62 in macrophages. **Lower panels**: Ratio of p62: β-actin. **C-E.** CRAMP^+/+^ and CRAMP^−/−^ macrophages were seeded in 35 mm dishes with 14 mm coverslips at 1× 10^6^/dish. The cells were stimulated with *FITC* labeled inactivated *E. coli* O22H8 (100 μl at 1 × 10^8^ CFU/ml) for 12 h. The samples were fixed with 4% neutral formalin for 5 min, stained with anti-mouse primary antibodies (1:100, anti-mouse LC3B, LAMP-1 and p62 antibodies) followed by a biotinylated anti-Ig secondary antibody (BD Biosciences) and streptavidin-PE. DAPI was used to stain nucleus. **C.** Reduced levels of LC3B proteins in CRAMP^−/−^ macrophages after stimulation with inactivated *E. coli* O22H8. Red: LC3B, Green: *E. coli*, Blue: DAPI. White arrow: LC3B, Yellow arrow: E. coli. Scale bar = 30 μm. **Right panel**: Quantitation of LC3B positive cells (%). \*\**P* < 0.01, two-tailed Student’s t test. **D.** Reduced levels of LAMP-1 proteins in CRAMP^−/−^ macrophages after stimulation with inactivated *E. coli* O22H8. Red: LAMP-1, Green: *E. coli*, Blue: DAPI. White arrow: LAMP-1, Yellow arrow: *E. coli*. Scale bar = 30 μm. **Right panel**: Quantitation of LAMP-1 positive cells (%). \*\**P* < 0.01, two-tailed Student’s t test. **E.** Increased levels of p62 proteins in CRAMP^−/−^ macrophages after stimulation with inactivated *E. coli* O22H8. Red: p62, Green: *E. coli*, Blue: DAPI. White arrow: p62, Yellow arrow: *E. coli*. Scale bar = 30 μm. **Right panel**: Quantitation of p62 positive cells (%). \*\**P* < 0.01, two-tailed Student’s t test.

Participation of CRAMP in the autophagy pathway in macrophages for *E. coli* elimination was further demonstrated by reduced fluorescence intensity of LC3B^+^ and LAMP-1^+^ (Lysosomal associated membrane protein 1) and increased fluorescence intensity of p62^+^ in *CRAMP*^*−/−*^ macrophages as compared to *CRAMP*^*+/+*^ counterparts after stimulation with inactivated *E. coli* for 12 h **(Fig. 5C-E)**. The bacteria showed reduced colocalization with LAMP-1 **(Fig. 5D)**, but increased colocalization with p62 **(Fig. 5E)** in *CRAMP*^*−/−*^ macrophages, indicating that CRAMP deficiency impaired degradation of bacteria conjugated with p62, resulting in retention of *E. coli* in the cells.

## Discussion

In this study, we elucidated previously uncharacterized macrophage effector mechanisms for elimination of phagocytosed *E. coli*. Viable *E. coli* infection or inactivated *E. coli* stimulation of mouse macrophages increase intracellular production and extracellular release of CRAMP by activation of NF-κB to trigger autophagy-dependent degradation of the bacteria ( as summarized in **Fig. 6)**. Interestingly, although both LPS and the chemotactic peptide fMLF are the products of *E. coli*, only LPS is able to up-regulate CRAMP expression in macrophages, indicating that TLR4 pathway promotes CRAMP expression and secretion similar to a previous report with mouse bone marrow-derived mast cells **(Li et al., 2009)**. In addition to TLR4, phagocytosed *E. coli* releases DNA to induce CRAMP/LL-37 also through interaction with TLR9 via activation of signal–regulated kinase (ERK) pathway **(Koon et al., 2011)**.

**Figure 6.**
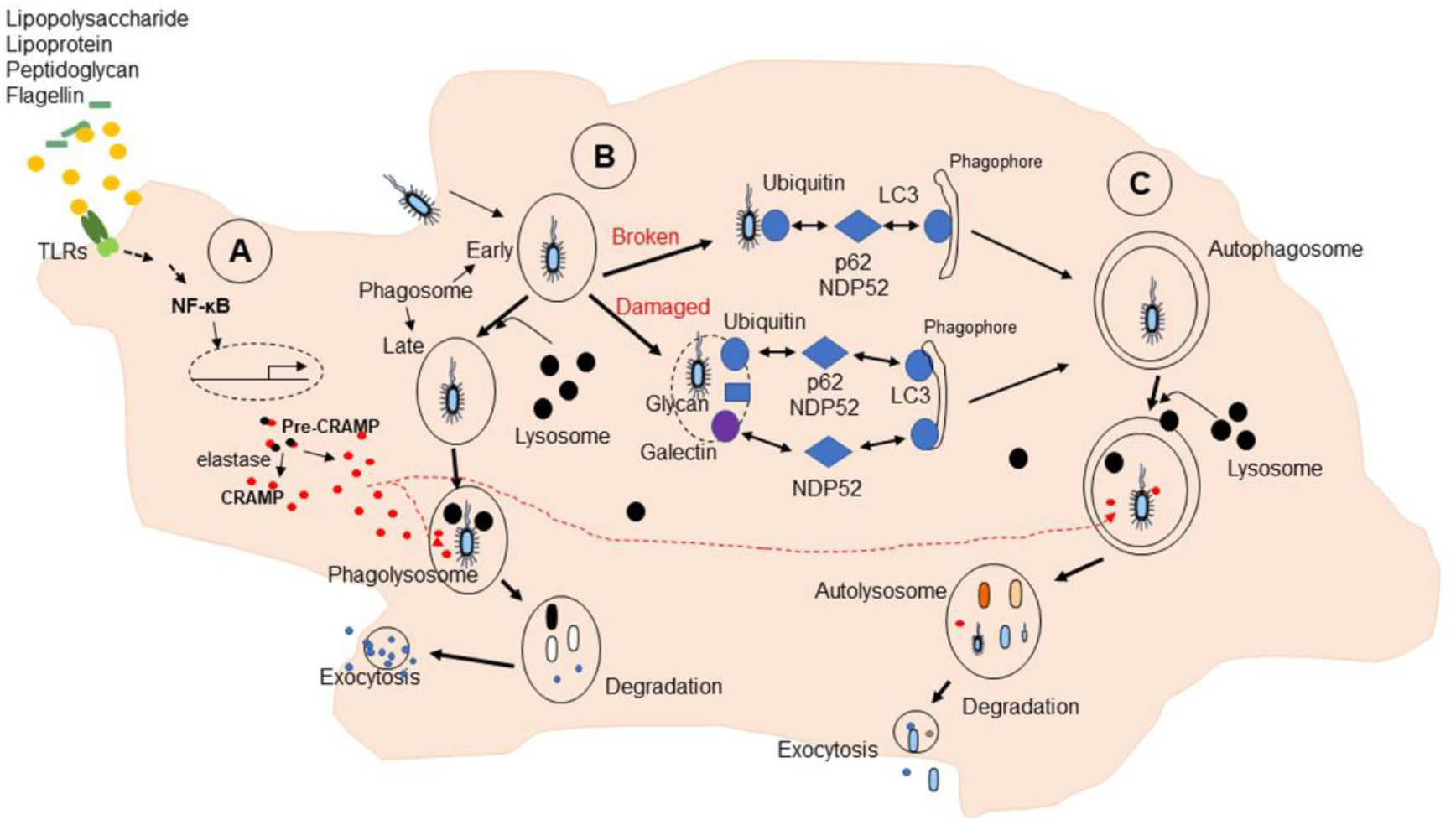
CRAMP-dependent autophagy to eliminate phagocytosed *E. coli* in macrophages. **A.** Up-regulation and activation of CRAMP. Soluble materials from E. coli stimulate TLRs-mediated signals to activate NF-κB, resulting in up-regulation of CRAMP expression. The pre-CRAMP is cleaved by elastase to form activated CRAMP. **B.** Selective capture and lysosomal degradation of cytosolic and vacuolar E. coli. E. coli phagocytosed in macrophages was incorporated a captured into phagosome, fused with lysosomes and degraded and. **C.** Autophagic pathway. Released naked E. coli from phagosomes or damaged phagosomes with E. coli in macrophages is captured by autophagosome either via ubiquitination. Autophagosomes fused with lysosomes in the form of autolysosomes are eventually degraded. NDP52: nuclear dot protein 52 kDa. P62: adaptor molecule p62 (also known as A170 or SQSTM1).

CRAMP/LL-37 plays an important role in intracellular bacterial killing by macrophages. It has been reported that (human?) macrophages kill *Mycobacterium tuberculosis* through vitamin-D3-induced CRAMP/LL-37 **(Yuk et al., 2009)**. Phenylbutyrate (PBA), an inducer of CRAMP/LL-37 in macrophages was able to restore LL-37 production originally suppressed by *M. tuberculosis* to enhance autophagy and intracellular bacterial killing by promoting colocalization of CRAMP/LL-37 and LC3-II in autophagosomes **(Rekha et al., 2015)**. The intracellular pathogen *Salmonella typhimurium* also increases macrophage production of CRAMP, which impairs *Salmonella* cell division, depending on intracellular elastase-like serine protease activity to proteolytically activate CRAMP **(Rosenberger et al., 2004)**. Ileal lesions in Crohn’s disease (CD) patients are abnormally colonized by pathogenic and invasive *E. coli* (AIEC). AIEC bacterial infection of macrophages induces the recruitment of autophagy machinery in the location of phagocytosis to limit intracellular AIEC replication. Impaired ATG16L1, IRGM or NOD2 expression in macrophages increases intracellular AIEC with enhanced secretion of IL-6 and TNF-*a* in response to infection. In contrast, forced induction of autophagy decreases the numbers of intra-macrophage AIEC and pro-inflammatory cytokine release **(Lapaquette et al., 2012)**. Our study observed that CRAMP inside macrophages plays an important role in sustaining the expression of autophagy-related proteins ATG5, LC3-II, LAMP-1 and p62 after phagocytosis of *E. coli*, as supported by the fact that *CRAMP*^*−/−*^ macrophages expressed lower levels of ATG5, LC3-II, LAMP-1 and p62 with impaired degradation of phagocytosed *E. coli* conjugated with p62 thereby culminating in accumulation of intracellular *E. coli*. These results further support the role of CRAMP-dependent autophagy in elimination of phagocytosed *E. coli* by macrophages.

CRAMP/LL-37 is expressed by various cells and tissues such as BM-derived myeloid cells (neutrophils, macrophages) and epithelial cells **(Zhang et al., 2019)**. CRAMP/LL-37 is stored in an intact form in the specific granules. During autophagy, CRAMP/LL-37 synthesized intracellularly is recruited to the autophagosomes **(Yuk et al., 2009)**. CRAMP/LL-37 contains both a conserved N-terminal cathelin-like region and a highly variable C-terminal region with bactericidal activity **(Cowland et al., 1995)**. The cathelin-like segment of antibacterial cationic proteins appear to be essential for subcellular trafficking through the synthetic apparatuses (ER, Golgi complex, and trans-Golgi network) **(Liu and Ganz, 1995)**. The release of active CRAMP/LL-37 from its precursor is mediated by proteinase 3 **(Sorensen et al., 2001)** or elastase **(Gudmundsson et al., 1996)**. We show that *E. coli* infection of macrophages increased CRAMP production. However, elastatinal blocks the capacity of macrophages to eliminate phagocytosed *E. coli*, suggesting that a critical concentration of active CRAMP is important for macrophage killing of intracellular *E. coli*.

Our current study showed that CRAMP deficiency is associated with reduced expression of autophagy-related proteins ATG5, LC3-II, LAMP-1 in macrophages after phagocytosis of *E. coli*. However, the changes in p62 are different. p62 (A170 or SQSTM1) is an accessory autophagy-targeting molecule that directs cytosolic proteins to autophagosomes, a process critical for the elimination of intracellular bacteria. p62 delivers specific cytosolic components, including ribosomal protein S30 (rpS30) and additional ubiquitinated proteins, to autophagic organelles to be processed into bactericidal products. In the absence of p62, the cells are unable to generate neo-antibacterial factors, resulting in non-functional autophagy despite maturation, thereby failing to effectively eliminate intracellular bacteria **(Ponpuak et al., 2010)**. The degradation of p62 is a widely used marker to monitor autophagic activity because p62 binds to LC3 and is selectively degraded by during autophagy **(Bjorkoy et al., 2005; Pankiv et al., 2007)**. In our study, CRAMP deficiency was shown to reduce the expression of p62 by macrophages at the early stages of *E. coli* infection, yet with significantly increased accumulation of *E. coli* colocalized with p62 at the later stage, indicating that autophagic process was impaired in the absence of CRAMP in association with bacterial retention.

We therefore disclosed a link between CRAMP and autophagy in macrophages, which assists the eradication of phagocytosed *E. coli*. These findings shed new light on the potential for consideration of developing autophagy-related therapies whereby innate immune responses are mobilized against infection and other diseases **(Levine and Kroemer, 2008)**, including IBD **(Haq et al., 2019; Kim et al., 2019; Larabi et al., 2020)** and neurodegenerative disorders **(Nixon, 2013)** with pathogenetic processes associated with defective autophagy activation.

## Materials and methods

### Mice

Myeloid cell-specific *CRAMP*^*−/−*^ (*LysMCre^+^CRAMP^F/F^*) mice were generated as described **(Chen et al., 2013b; Yoshimura et al., 2018).** Mice used in the experiments were 8–12 weeks old and were allowed free access to standard laboratory chow/tap water. All animals were housed in an air-conditioned room with controlled temperature (22 ± 1°C), humidity (65–70%), and day/night cycle (12 h light, 12 h dark). All animal procedures were governed by the US NIH Guide for the Care and Use of Laboratory Animals (National Academies Press, 2011) and were approved by the Animal Care and Use Committee of the NCI-Frederick, National Institutes of Health.

### Generation of CRAMP^+/+^ control and CRAMP^−/−^ macrophages

BM was flushed from the femurs of euthanized mice with PBS as described **(Chen et al., 2013a)**. Red cells were lysed with ACK Lysing Buffer (Cambrex Bio Science, MD). Cell suspension was centrifuged for 10 min at 1200 rpm and the pellet was gently resuspended in Dulbecco’s modified essential medium (DMEM; Gibco Invitrogen) supplemented with 2 mM L-glutamine (Gibco Invitrogen), 10 mM Hepes (Gibco Invitrogen), 1 mM sodium pyruvate (Gibco Invitrogen), 10% heat-inactivated FBS (Gibco Invitrogen) and 50 ng/ml M-CSF. To remove fibroblasts, the cells were cultured in tissue culture dishes (Corning Inc. NY) at 37°C in a 5% CO2 overnight. The non‐adherent cells were collected, centrifuged and re-cultured in tissue culture dishes (1×10^6^ cells/ml) with addition of DMEM with 50 ng/ml M-CSF for 3 days. The medium was replaced on day 7 and fully differentiated macrophages were harvested. *CRAMP*^*+/+*^ macrophages were generated from BM of control (*LysMCre*^*−*^*CRAMP*^*F/F*^) mice (referred to as control cells) and *CRAMP*^*−/−*^ macrophages were generated from BM of Myeloid cell-specific *CRAMP*^−/−^ (*LysMCre*^*+*^*CRAMP*^*F/F*^) mice.

### Isolation of fecal E. coli

The fecal *E. coli* isolated from naïve mice was aerobically cultured on Violet red bile lactose (VRBL) agar for 24 h. Single bacterial colonies were identified as *E. coli* O22H8 by complete genome sequencing. *E. coli* identified was cultured in LB agar at 37°C, 180 RMP for 24 h, then determined for concentrations at OD_600nm_ = 0.4 corresponding to ~2 × 10^8^ colony forming unit (CFU)/ml). *E. coli* suspension was aliquoted in 1 ml volumes and stored at −80°C for future use. When necessary, live or heat-inactivated *E. coli* was labeled with FITC (Isomer I, Sigma) following the manufacturer recommended procedures.

### Detection of CRAMP produced by CRAMP^+/+^ macrophages

BM-derived *CRAMP*^*+/+*^ control macrophages were seeded at 1.5 × 10^5^ cells/well in 96 well plates. Live or heat inactivated *E. coli* was added into the wells at 1× 10^7^ CFU/well (MOI=100). After culture at 37°C in 5% CO2, cell supernatant was harvested at different time points to measure CRAMP concentration with ELISA using a Mouse CRAMP ELISA Kit (MyBioSource, CA).

### In vitro killing of E. coli by CRAMP

*E. coli* was diluted at 5 × 10^4^ in 100 μl/well on 96-well plates followed by culture with various concentrations (0.01 - 100 μg/ml) of synthetic murine CRAMP (Hycult Biotech) at 37°C for 2 h. The bacterial suspension was then serially diluted with PBS and plated on nutrient agar plates at 37°C for 24 h. The number of *E. coli* treated with CRAMP was quantitated and expressed as the percentage of the number of untreated bacteria as a control.

### Killing of phagocytosed E. coli by macrophages

BM-derived *CRAMP*^*+/+*^ control and *CRAMP*^*−/−*^ macrophages seeded in 35 mm dish with 14 mm coverslips in the bottom (MatTeck Corporation, MA) were infected with *E. coli* isolated at a multiplicity of infection of 5 live bacteria per cell at 37°C in DMEM supplemented with 10% FCS in the presence of M-CSF (50 ug/ml) without antibiotics. The cells were fixed at 0, 4, and 20 h then stained with SYTO 9 (ThermoFisher), a Gram stain Kit (abcam) or an anti-*E. coli* antibody (Ab) and Gram strain kit.

### Elimination of phagocytosed E. coli by macrophages

BM-derived *CRAMP*^*+/+*^ control and *CRAMP*^*−/−*^ macrophages were seeded in 35 mm dish with 14 mm coverslips in the bottom at 1× 10^6^ cells/dish and co-cultured with FITC-labeled inactivated *E. coli* at a multiplicity of 10 bacteria per cell at 37°C in DMEM supplemented with 10% FCS in the presence of M-CSF (50 ug/ml). The cells were fixed at 0, 4, 6 and 24 h then stained with DAPI for nuclei. The percentages (%) of macrophages containing phagocytosed inactivated *E. coli* and the number of phagocytosed inactivated *E. coli* in single macrophage at indicated time points were measured.

### Immunofluorescence

BM-derived *CRAMP*^*+/+*^ control and *CRAMP*^*−/−*^ macrophages were seeded in 35 mm dish with 14 mm coverslips in the bottom at 1× 10^6^ cells/dish and co-cultured with FITC-labeled inactivated *E. coli* at a multiplicity of 10 bacteria per cell at 37°C in DMEM supplemented with 10% FCS in the presence of M-CSF (50 ug/ml) for 12 h. The cells were fixed with 4% neutrally buffered formalin for 5 min and stained with anti-mouse primary antibodies (1:100, anti-mouse LC3B, LAMP-1 and p62 antibodies) followed by a biotinylated anti-Ig secondary antibody (BD Biosciences) and streptavidin-PE. DAPI was used to stain nucleus. 4-8 viewing fields from each slide were captured under fluorescent microscopy.

### Western immunoblotting

BM-derived *CRAMP*^*+/+*^ control and *CRAMP*^*−/−*^ macrophages or CT26 mouse colon epithelial cell line (ATCC) grown in 60-mm dishes to sub-confluency were cultured for 3 h in FCS-free media. After treatment with inactivated *E. coli*, the cells were lysed with 1× SDS sample buffer (62.5 mM Tris–HCl (pH 6.8), 2% SDS, 10% glycerol, and 50 mM dithiothreitol), then sonicated for 15 s and heated at 100°C for 5 min. Cell lysate was centrifuged at 12,000 rpm (4°C) for 5 min, and protein concentrations of the supernatants were measured by DC Protein Assay (Bio-Rad). The lysates with titrated proteins were electrophoresed on 10% SDS-PAGE precast gels (Invitrogen) then transferred onto ImmunoBlot polyvinylidene membranes (Bio-Rad), which were blocked with 5% nonfat milk. Phosphorylated IκB-α were detected using phospho-specific Abs according to the manufacturer’s instructions. After incubation of the membranes with a horseradish peroxidase-conjugated secondary Ab, protein bands were detected with a Super Signal Chemiluminescent Substrate (Pierce) and the images were quantitated using a G-BOX GeneSnap system (SYNGENE). For detection of total IκB-α, β-actin, ATG-5, LC3B, p62 and CRAMP, the membranes were stripped with Restore Western Blot Stripping Buffer (Pierce) followed by incubation with specific Abs.

### Statistics

All experiments were performed at least three times with three replicate samples. Statistical analysis was performed using GraphPad Prism by two-tailed Student’s t test or 1-way ANOVA with Kruskal-Wallis Test. Data with error bars represent mean ± SEM and *P* value*s* less than 0.05 (*P* < 0.05) were considered statistically significant.

## Acknowledgements

This project was funded in part by federal funds from the National Cancer Institute, National Institutes of Health, under Contract No. HHSN261200800001E and was supported in part by the Intramural Research Program of the NCI, NIH, and by the fund from LCIM, CCR, NCI-Frederick. The authors thank Ms. Cheri A. Rhoderick for secretarial assistance. The authors also thank Drs. Amiran K. Dzutsev, and John McCulloch for insightful advice and sequencing of *E. coli*.

## Author Contributions

Conceptualization, K.C. and J.M.W.; Methodology, K.C. W.G. and J.M.W; Investigation, K.C., T.Y., J.H., W.G, and C.T.; Intellectual inputting, G.T.; Writing-original draft, K.C.; Reviewing & Editing, J.M.W.; Supervision, J.M.W.

